# Regulation of Adult Zebrafish Retinal Regeneration by Lamβ1b-Chain-Containing Laminins

**DOI:** 10.1101/2025.06.24.661136

**Authors:** Dmitri Serjanov, James Twist, Haoyuan Jiang, David R. Hyde

## Abstract

Retinal degenerative diseases are a major cause of blindness in humans that often result in permanent and progressive loss of vision. Unlike humans, zebrafish possess the remarkable ability to regenerate lost retinal neurons through Müller glia (MG) reprogramming and asymmetric cell division to produce multipotent retinal progenitor cells (RPCs). While most studies on the molecular mechanisms underlying this regeneration process have focused on intracellular mechanisms, the role of the microenvironment surrounding retinal cells, the extracellular matrix (ECM), has been understudied. Laminins are heterotrimeric glycoproteins, are principal components of the ECM basement membrane, and play important roles in vertebrate retinal development. Here, we examine the role of β1b chain-containing laminins in the regenerative response of the zebrafish retina. We found that the zebrafish *lamb1b* gene is differentially expressed during MG reprogramming and MG and NPC proliferation during retinal regeneration. Further, we found that β1b-containing laminins play important roles in regulating MG and NPC proliferation and neuroprotection of photoreceptors in light-damaged zebrafish retinas. Finally, Lamβ1b plays an important role in regulating the expression of integrin receptors and other laminin genes during the regeneration response. Taken together, Lamβ1b, and likely other ECM components, play a critical role in the MG-dependent neuronal regeneration response in the zebrafish retina.

## INTRODUCTION

Unlike humans, zebrafish possess an extensive regenerative capacity, including the ability to regenerate all cell types in damaged retinas (Vihtelic and Hyde, 2000; Lahne et al., 2020). Following retinal damage, zebrafish Müller glia (MG) respond by entering a transient gliotic state similar to mammals (Thomas et al., 2016; Hoang et al., 2020). However, the zebrafish MG then reprogram to a transcriptional state similar to a late retinal progenitor cell (Lyu et al., 2023). These reprogrammed MG undergo asymmetric cell division to produce a Müller glia-derived Neuronal Progenitor Cell (MGPC), which continues to proliferate and migrate to the location of damaged tissue, where they differentiate into all missing retinal cell types (Vihtelic and Hyde, 2000; Cameron et al., 2005; Fausett and Goldman, 2006; Vihtelic et al., 2006; Bernardos et al., 2007; Kassen et al., 2007). There are many MG genes/proteins known to regulate this regeneration response (Lahne et al., 2020), as well as several extrinsic proteins. For example, TNF*α* is released from dying photoreceptors after 16 hours of light-induced retinal damage and is required for MG asymmetric cell division (Nelson et al., 2013). Additionally, several groups have studied the role of microglia in regulating MG-dependent retinal regeneration (White et al., 2017; Mitchell et al., 2019; Dyck et al., 2021; Iribarne and Hyde, 2022; Conedera et al., 2023; Lu and Hyde, 2024). Thus, MG reprogramming and reentry into the cell cycle requires signals extrinsic to the MG.

One potential extrinsic MG signal that remains significantly understudied in retinal regeneration is the extracellular matrix (ECM). The ECM is a dynamic non-cellular structure composed of a diverse array of proteins that acts as a physical scaffold and regulates developmental and homeostatic signaling pathways (Yurchenco, 2011; Varshney et al., 2015). Various cell types contribute to the composition of retinal ECM, including the MG (Libby et al., 2000b; Halfter et al., 2008). Within the retina, the ECM is present in several forms and locations: the inner limiting membrane (ILM) at the vitreal surface, outer and inner plexiform regions, the inter-photoreceptor matrix (IPM), and Bruch’s membrane (Reinhard et al., 2015). Both ILM and Bruch’s membrane are basement membranes (BMs), a specialized sheet-like form of ECM that serves as a physical barrier between the retina and the surrounding tissues of the eye.

During vertebrate eye development, extrinsic environmental cues, such as those from the ECM, are necessary for cell-fate determination, proliferation, differentiation, and migration (Varshney et al., 2015). ECM molecules indirectly control developmental processes through their ability to alter the diffusion rates and local concentrations of soluble molecules in the retina by sequestering the molecule (Libby et al., 2000a). The ECM can also directly affect eye development through ILM-cell signaling interactions that are critical for achieving proper retinal composition (Reinhard et al., 2015). These interactions are facilitated by an array of cell-surface receptors, such as integrins, that directly link the ECM to the internal cytoskeleton of retinal cells (Yurchenco, 2011). Because regeneration is similar to retinal development (Lyu et al., 2023) and ECM components are critical for development, it is likely that the ECM will also be involved in retinal regeneration.

One important component of the ECM is laminins, a family of heterotrimeric glycoproteins that are principal components of the ECM and indispensable for BM assembly (Smyth et al., 1999; Tunggal et al., 2000; Miner and Yurchenco, 2004). Laminin trimers are composed of an *α*, *β*, and *γ* chain, assembled through *α*-helical coiled-coils (Yurchenco, 2011). There are 5 *α*, 3 *β*, and 3 *γ* vertebrate isoforms, which can combine into 16 distinct laminin trimers (Domogatskaya et al., 2012). Although the *α* chain directly mediates laminin-cell binding, the β and *γ* chains can also modulate these interactions (Domogatskaya et al., 2012). Additionally, the β chain contains an LN domain at the N-terminus, which facilitates self-polymerization into a lattice network, forming the BM foundation that anchors onto the cell surface (Cheng et al., 1997; Durbeej, 2009). Laminins are known to mediate cell adhesion, proliferation, migration, differentiation, and apoptosis by binding to cells, initiating a signaling cascade, and eventually altering gene expression (Xu et al., 2010). However, the role of laminins in retinal regeneration is currently unknown.

In this study, we identified laminin beta 1b (*lamb1b*), a critical component for BM formation (Edwards and Lefebvre, 2013), as being significantly upregulated during regeneration following constant bright light damage. Additionally, we show that Lamβ1b serves a neuroprotective role for photoreceptors during constant light treatment and is also a necessary component of the regenerative response in the zebrafish retina. Specifically, β1 chain containing laminins (CCLs) appear to regulate MG and MGPC proliferation and may regulate the expression of other ECM components, influencing the overall composition of the retina ECM during regeneration. Considering the crucial role of β1 CCLs in retinal development, these findings suggest a similar role in regeneration.

## MATERIALS AND METHODS

### Zebrafish husbandry

Adult female and male zebrafish (*Danio rerio*), *albino* and Tg(*gfap:EGFP*)^nt11^, were bred and maintained at 26.5°C under normal light conditions (14h light:10h dark) in the Freimann Life Science Center (FLSC) Zebrafish Facility at the University of Notre Dame in accordance with the FLSC standard operating policies and previously described procedures (Vihtelic and Hyde, 2000). Transgenic Tg(*gfap:EGFP*)^nt11^ zebrafish express enhanced green fluorescent protein (eGFP) using the *gfap* promoter in Müller glia (MG; (Kassen et al., 2007). All the zebrafish used in this study were adults between 6 and 18 months old and 3-5 cm in length. Prior to any experiments, the fish were anesthetized or euthanized using either 1:1000 or 1:500 2-phenoxyethanol (2-PE; 77699; Sigma-Aldrich, St. Louis, MO) in system water, respectively. All animal care protocols were approved by the University of Notre Dame Animal Care and Use Committee and are in compliance with the Association of Research in Vision and Ophthalmology for the use of animals in vision research.

### Light treatment protocol

Phototoxic treatment that causes photoreceptor death was used to initiate a regenerative response in the zebrafish retina (Vihtelic and Hyde, 2000). All fish subjected to light treatment were adapted in constant darkness for two weeks and then placed immediately adjacent to intense fluorescent lights for the desired amount of time, which varied between 6 hours and 96 hours (Table 1). The temperature of the fish tanks was regularly monitored and maintained at 34°C. In some cases following light treatment, the fish were returned to standard light:dark conditions for recovery and observation.

**Table 1-.**
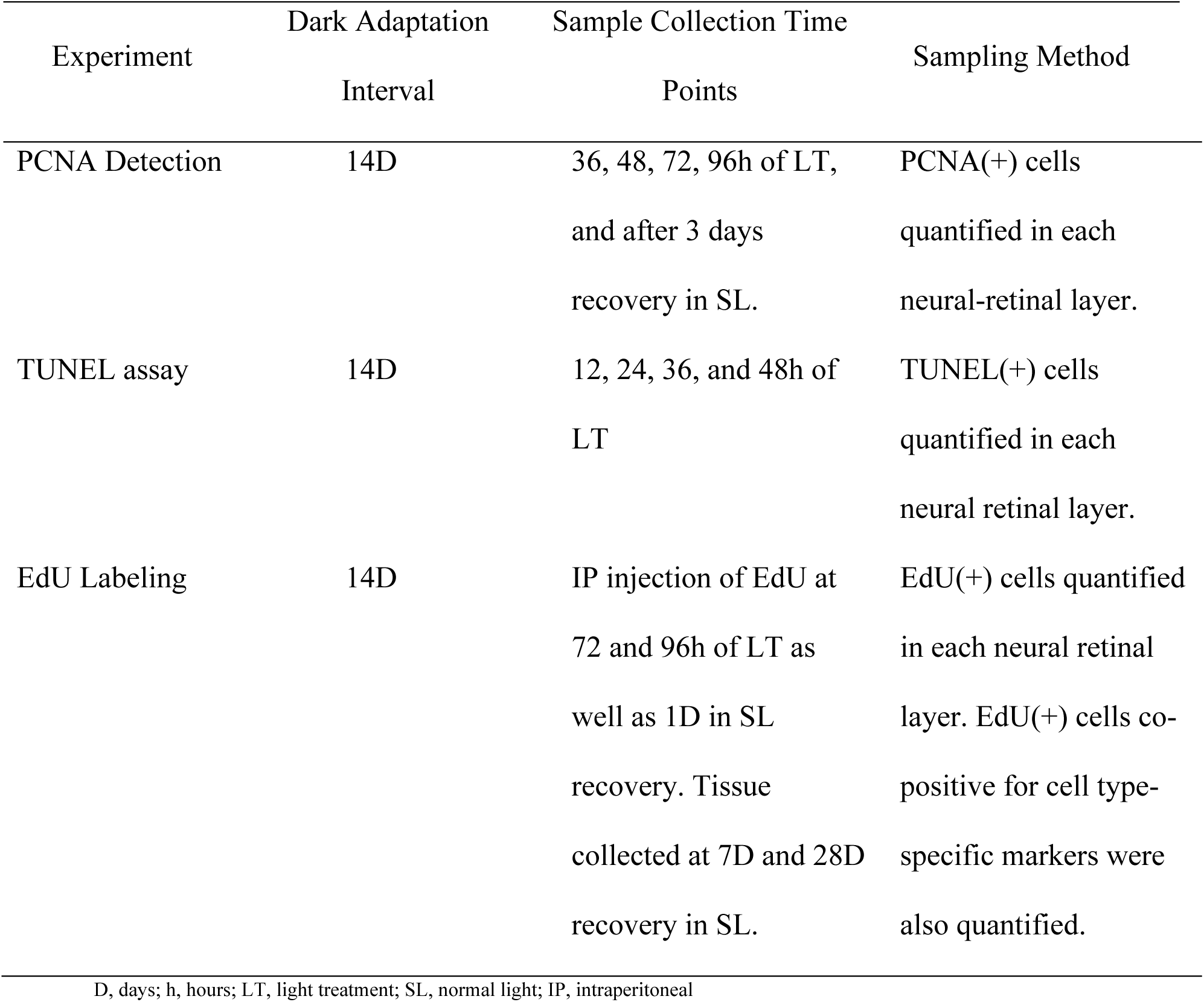
Experimental Protocols.

### *In vivo* morpholino-mediated gene knockdown

Prior to constant light treatment, fish were anesthetized using 1:1000 2-phenoxyethanol and intravitreally injected with a fluorescent lissamine-tagged morpholino (Gene Tools, LLC) that is complementary to *lamb1b* mRNA, followed by immediate electroporation into the retina using a standard procedure developed by the Hyde lab (Thummel et al., 2011). Using Dumont #5 forceps, the outer cornea of the left eye was removed and an incision was made through the inner cornea in the ventral-caudal aspect of the pupillary margin using a #11 scalpel. Using a 2.5µL Hamilton syringe, 0.2-0.3µL of 1 mM morpholino was injected into the posterior segment, displacing some vitreous. Electrodes were used to prolapse the eye out of the socket, and the negative electrode was positioned behind the dorsal part of the eye. Electroporation was performed using electrode tweezers connected to a CUY21EDIT Square Wave Electroporator (Nepa Gene Company Ltd.) that delivers two 50 msec pulses of electricity, separated by 1 second, at 0.75 Volts and 0.05 Amperes. Electroporation drove the morpholinos into the dorsal retinal cells so they may basepair with complementary mRNAs and suppress translation of the encoded protein. Control morpholinos included a Standard Control that is not complementary to any known sequence in the zebrafish genome (Gene Tools, LLC) and a *pcna* morpholino that represses Müller glia cell proliferation. Successful electroporation was confirmed by the presence of lissamine fluorescence within the retinal cells using confocal microscopy. The right eyes were used as uninjected controls. The morpholinos used in this study are listed in Table 2 and were stored at room temperature in 1mM stocks.

**Table 2-.**
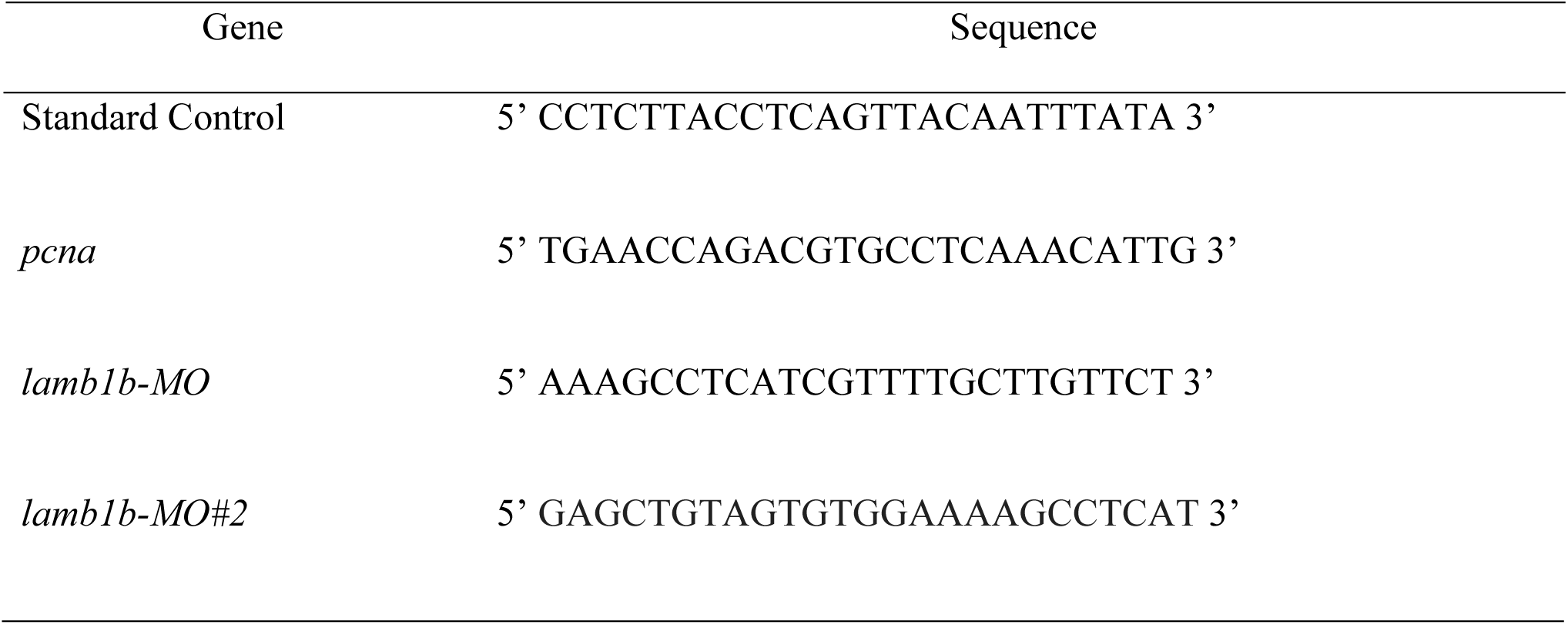
Morpholino Sequences.

### Cryosectioning

Zebrafish eyes were enucleated using #5/45 Dumont forceps. The eyes were then floated in 1x Phosphate Buffered Saline (PBS) solution and a circular opening, approximately the size of the pupil, was made in the cornea using micro-scissors, creating an “eye cup.” The eye cups were fixed at room temperature (RT) in 4% paraformaldehyde, using a pH shift technique that is optimized to preserve cytoskeletal structures (see Tissue fixation). After washing the eyes in 1x PBS, the tissue was dehydrated in a 15% sucrose solution at 4°C overnight. The eyes were then transferred to a solution of two parts PolyFreeze Tissue Freezing Medium (TFM) (Sigma-Aldrich) to one part 30% sucrose for another 24 hours at 4°C. Finally, the eyes were arranged in 100% TFM and stored at −80°C until cryosectioning. Using a Thermo Scientific Microm HM550 Cryostat, the frozen tissue blocks were sectioned at −22°C, producing 14µM serial retinal slices that included the dorsal and ventral retina, as well as the optic nerve. The retinal sections were collected on glass slides, dried for 30 minutes on a slide warmer at 50°C, and stored at −80°C for future immunohistochemical staining.

### Tissue fixation

We used a pH shift fixation technique to preserve cytoskeletal structure and to avoid the cellular swelling that is typically associated with standard paraformaldehyde (PFA) fixation. Here, we used two solutions of 4% PFA at a pH of 7 and 11, which are referred to as “Solution I” and “Solution II,” respectively. Eyes were fixed in Solution I (80mM HEPES, 2mM EGTA, 2mM MgCl_2_, 4% PFA, pH 6.8-7.2) at room temperature for 5 minutes, then transferred to Solution II (100mM Na_2_B_4_O_7,_ 1mM MgCl_2_, 4% PFA, pH 11) for a longer 30-minute fixation on a rocker at room temperature.

### Immunohistochemistry

Immunohistochemistry of retinal sections was performed similar to what was previously described (Lyu et al., 2023). Glass slides containing retinal sections were removed from −80°C storage and dried on a slide warmer for 30 minutes at 50°C. The slides were then washed in a Coplin jar for 3×10 minutes in 1xPBS to remove residual Tissue Freezing Medium. The slides were incubated in a blocking solution of 5% Normal Goat Serum (Vector Laboratories, Inc.), 0.3% Triton X-100, and 94.7% 1xPBS for 30 minutes at RT and then incubated in primary antibodies that were diluted to various concentrations in antibody dilution solution of 5% Normal Goat Serum (Vector Laboratories, Inc.), 0.01% Triton X-100, and 95% 1xPBS overnight at RT in a humid chamber. The slides were then washed for 3×10 minutes in 1xPBS at RT. Secondary antibodies in various concentrations were diluted in antibody dilution solution and applied to the slides for 4 hours at RT. The slides were then washed in 1xPBS three times for 10 minutes each at RT. Finally, glass coverslips were mounted on the slides using ProLong^TM^ Gold antifade reagent (Invitrogen). When anti-PCNA primary antibodies were used, an extra heat-induced antigen retrieval step was performed to facilitate the antibody’s ability to detect its epitope. Prior to blocking, the slides were immersed in sodium citrate solution (10mM Sodium Citrate, 0.05% Tween 20, pH 6) and heated for 25 seconds in a microwave at maximum power. The slides were heated for another 7 minutes using the microwave at 10% power in periodic 20-25 second bursts. Care was taken to heat the sodium citrate solution without bringing it to a boil. The slides were allowed to cool for 30 minutes at RT and washed 2×5 minutes in 1xPBS before proceeding to the blocking step. The remainder of the staining process proceeded as previously described. All primary and secondary antibodies used in this study are listed in Table 4.

**Table 4-.**
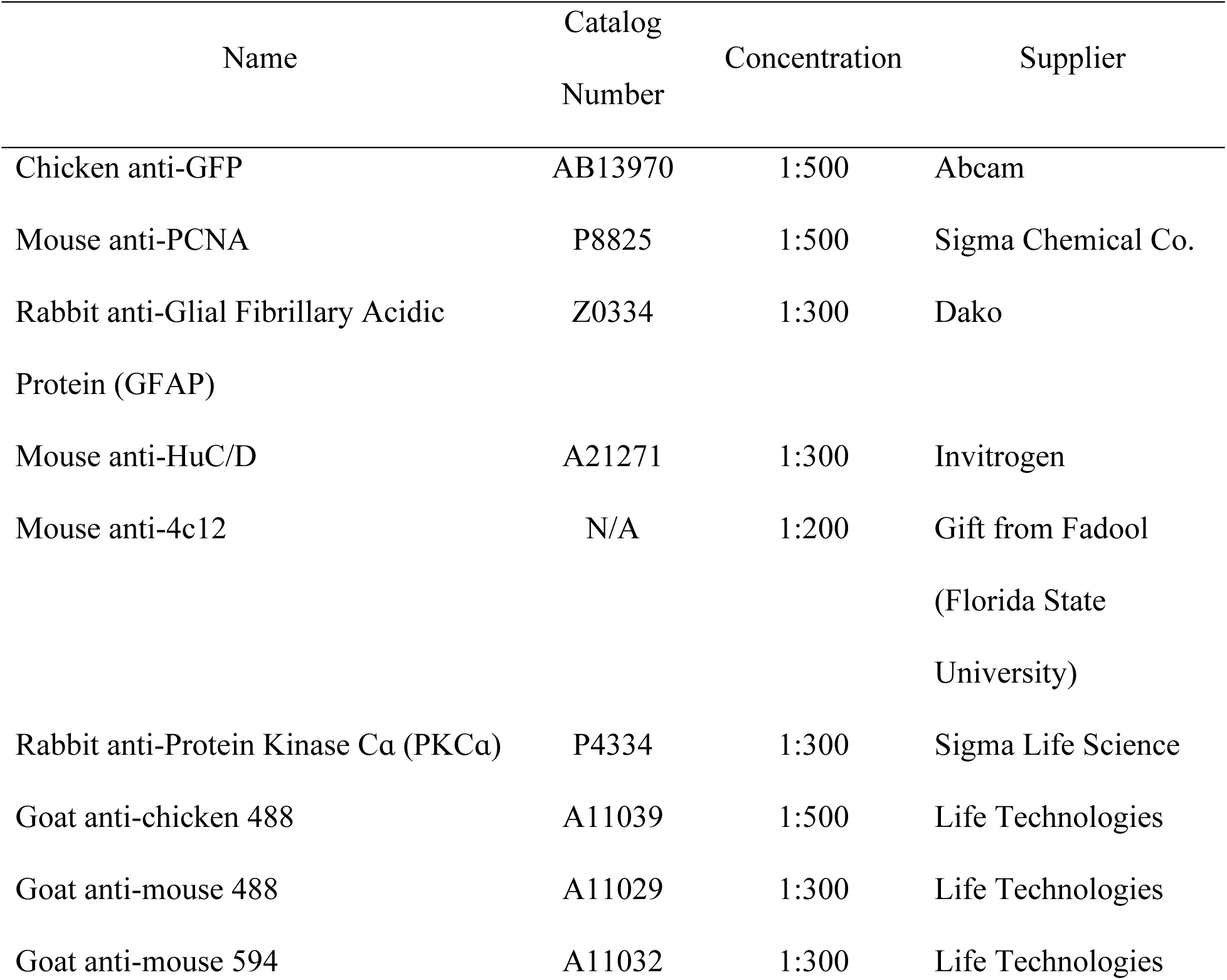
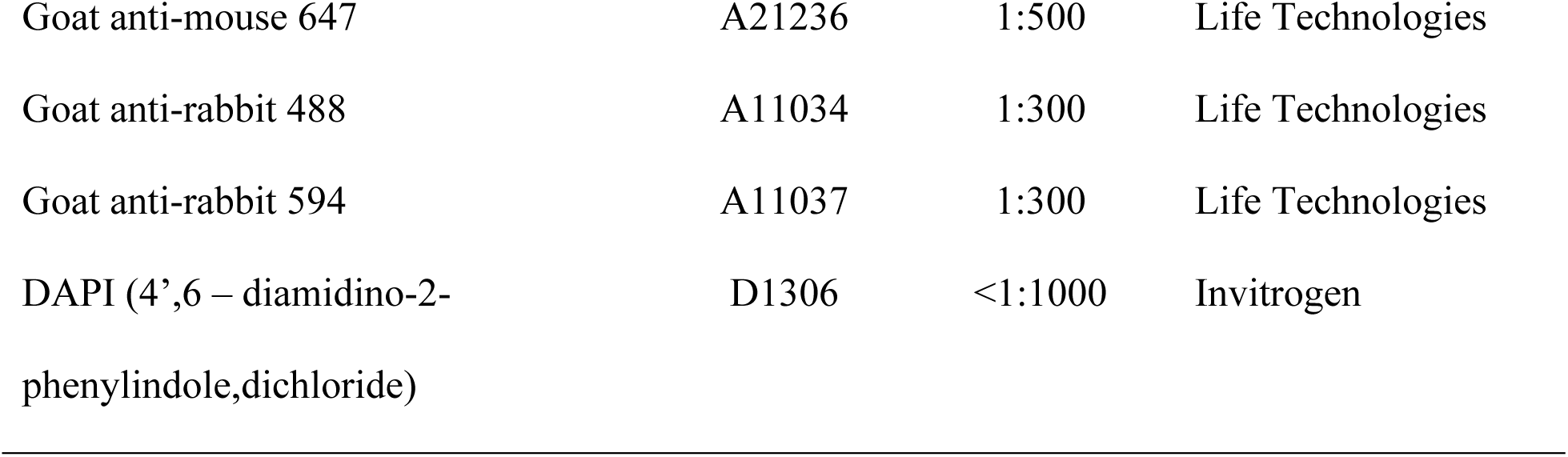
Primary and Secondary Antibodies.

| Name | Catalog<br>Number | Concentration | Supplier |
| --- | --- | --- | --- |
| Chicken anti-GFP | AB13970 | 1:500 | Abcam |
| Mouse anti-PCNA | P8825 | 1:500 | Sigma Chemical Co. |
| Rabbit anti-Glial Fibrillary Acidic<br>Protein (GFAP) | Z0334 | 1:300 | Dako |
| Mouse anti-HuC/D | A21271 | 1:300 | Invitrogen |
| Mouse anti-4c12 | N/A | 1:200 | Gift from Fadool<br>(Florida State<br>University) |
| Rabbit anti-Protein Kinase Cα (PKCα) | P4334 | 1:300 | Sigma Life Science |
| Goat anti-chicken 488 | A11039 | 1:500 | Life Technologies |
| Goat anti-mouse 488 | A11029 | 1:300 | Life Technologies |
| Goat anti-mouse 594 | A11032 | 1:300 | Life Technologies |
| Goat anti-mouse 647 | A21236 | 1:500 | Life Technologies |
| Goat anti-rabbit 488 | A11034 | 1:300 | Life Technologies |
| Goat anti-rabbit 594 | A11037 | 1:300 | Life Technologies |
| DAPI (4',6 – diamidino-2-phenylindole,dichloride) | D1306 | <1:1000 | Invitrogen |

### TUNEL assay

Detection of double-stranded DNA breaks, indicative of apoptosis, was performed on retinal sections by terminal deoxynucleotidyl transferase dUTP nick-end labeling (TUNEL) as previously described (Vihtelic et al., 2006). Glass slides containing fixed retinal sections were prepared as above. The slides were washed, blocked, and incubated in both primary and secondary antibody solutions as described above. Following secondary antibody incubation, the slides were washed for 5 minutes in 1xPBS and fixed by incubation in a 4% PFA solution at RT. Unless otherwise stated, all subsequent washes were performed using 1xPBS in a Coplin jar at RT. Next, the slides were incubated in a 1:1000 solution of Proteinase K (Clontech) in 1xPBS at RT. After two 10-minute washes, antigen retrieval was performed by immersing the slides in a chilled sodium citrate solution (3% Sodium Citrate, 0.1% Triton) at −20°C for 5 minutes. The slides were washed 2 times for 2 minutes each before incubating in an equilibration buffer (Clontech) at RT. Next, the slides were incubated in a label mix (96% equilibration buffer, Clontech; 2% dNTP-biotin, R&D Systems; 2% TdT transferase, Takara) for 2 hours at 37°C. After the DNA labeling reaction, the slides were washed for 15 minutes in 2x saline sodium citrate buffer (SSC, Takara) at RT. Finally, the slides were washed briefly two times for 2 minutes each, and then incubated in 1:200 Streptavidin, Alexa Fluor 647 conjugate (Invitrogen), washed 2×5 minutes, and mounted in ProLong^TM^ Gold antifade reagent (Invitrogen) and a coverslip (Avantor). Finally, labeled apoptotic cells were visualized using a Nikon A1 inverted confocal microscope.

### EdU injections and detection

EdU (5-Ethynyl-2′-deoxyuridine) was incorporated into the DNA of proliferating cells as previously described (Lyu et al., 2023). First, the fish are anesthetized using 1:1000 2-phenoxyethanol and immobilized under a paper towel in a petri dish. Using an insulin syringe with a 30-gauge needle, approximately 0.05 mL of 1xEdU (1mg/ml) was intraperitoneally injected into each zebrafish. These injections were performed after either 72 hours or 96 hours of constant light treatment. In some experiments, EdU injections also took place at 1 day of recovery under normal light conditions (1 day following 96 hours of constant light). The different timepoints when these single EdU injections occurred (see Table 1), allow for increased temporal resolution in the detection of dividing cells. To detect EdU in retinal sections affixed to glass slides, slides were incubated in blocking solution (see Immunohistochemistry). Then, the slides were incubated at RT for 30 minutes per manufacturer’s instructions (Invitrogen), using Alexa Fluor 647 Azide (Invitrogen, Catalog Number A10277) for visualization. The slides were washed in 1xPBS at RT 2×5 minutes at RT before proceeding with immunohistochemistry.

### Confocal microscopy and cellular quantification

A Nikon A1R HD inverted confocal system was used to capture z-stack serial optical sections of retinal slices affixed to a glass slide. All images were captured at the University of Notre Dame Integrated Imaging Facility. In general, a 40x oil objective was used to capture the z-stacks of approximately 10µM of dorsal retinal tissue. Immunolabeled cells were counted in maximum projection z-stacks (PCNA+, EdU+, PKC*α*+, HuC/D+, cones) as well as single-slices (GFAP+, 4c12+). After normalization of cells per 100µM, a one-way ANOVA and Tukey’s post-hoc test were used to determine statistical significance between the control retina, *pcna* morphants, and *lamb1b* morphants for all experiments.

### RNA extraction

To dissect the retinal tissue from the rest of the eye, we first made an “eye cup” using the methods described under “*Cryosectioning*.” Importantly, this procedure was performed without any fixation and in a small petri dish on ice with the aid of a light microscope. Using micro-scissors and #5 Dumont forceps to stabilize the eye, a cut was made along the cranial-caudal axis to separate the eye into ventral and dorsal halves because the dorsal retina is most affected by the constant light damage (Vihtelic et al., 2006). Once two halves of the eye were created, the sclera and choroid layers were carefully dissected away from the retina using #5 Dumont forceps to keep the delicate retinal tissue intact. Finally, the retinal pigment epithelium was scraped away from each of the retinas and immediately frozen in 1.5mL sterile Eppendorf tubes on dry ice. The dorsal and ventral halves of the eye were stored separately at −80°C until future use. In a fume hood, 500µL of TRIzol (Ambion) was added to a 1.5µL Eppendorf tube with approximately 10 dorsal retinas. The tissue was homogenized in TRIzol solution and the tubes were placed on a rocker for 5 minutes at RT. Next, 100µL of chloroform was added in the fume hood at RT, the tubes were mixed by shaking, and placed on the rocker for 3 minutes at RT. The tubes were centrifuged for 15 minutes at 12,000 rpm at 4°C to separate the aqueous and phenol phases. The aqueous phase was transferred to a new sterile 1.5µL Eppendorf tube by pipetting from the top down to avoid pipetting any of the DNA, which is found at the interface between the aqueous and phenol phases. The DNA and the phenol phase, which contains proteins, were stored in −80°C for future use. A volume of 1mL of 100% isopropanol was added to the aqueous phase containing the RNA, and the solution was mixed by shaking and vortexing. After letting the solution rest for 10 minutes at RT, it was centrifuged for 10 minutes at 12,000 rpm at 4°C. The supernatant was discarded and the RNA pellet was dissolved by mixing in 500µL of RNAse-free dH_2_O (Invitrogen). Next, 500µL of isopropanol, 50µL of 3M NaAc (pH 5.5), and 10µL of 1mg/mL glycogen were added to the solution, mixed and left to rest at RT for 20 minutes. After centrifuging for another 10 minutes at 12,000 rpm at 4°C, the supernatant was discarded, and 500µL of ice-cold 75% ethanol was added to the pellet and gently mixed. The tube was centrifuged again for 5 minutes at 12,000 rpm at 4°C. We discarded the supernatant, and the pellet was ethanol washed again. The pellet was left to air-dry for 15 minutes in a fume hood, and then resuspended in 25µL of RNAse-free dH_2_O. Following concentration and purity measurements using a NanoDrop, RNA samples were stored at −80°C until future use.

### cDNA synthesis

Synthesis of cDNA was performed using a Peltier Thermal Cycler-200 (MJ Research). Reactions were prepared by combining RNA samples with qScript cDNA SuperMix (QuantaBio) following the manufacturer’s protocols. The thermocycler was programmed for 5 minutes at 25°C, 30 minutes at 42°C, followed by 5 minutes at 85°C. DNA concentration was measured using a NanoDrop.

### Quantitative real-time polymerase chain reaction (qRT-PCR)

Reactions for qRT-PCR were prepared by combining cDNA samples with SYBR Green SuperMix (Quantabio) along with the appropriate primer pairs, according to manufacturer’s protocols. A StepOnePlus Real-Time PCR System (Applied Biosystems) was programmed to a 10-minute holding stage at 95°C, followed by a cycling stage of 40 cycles of 15 seconds at 95°C and 1 minute at 60°C. Melting curves were measured at 95°C for 15 seconds, followed by 60°C for 1 minute, and 95°C for 15 seconds. All forward and reverse primer sequences used in this study are listed in Table 5. *18s* was used as a reference gene.

**Table 5-.**
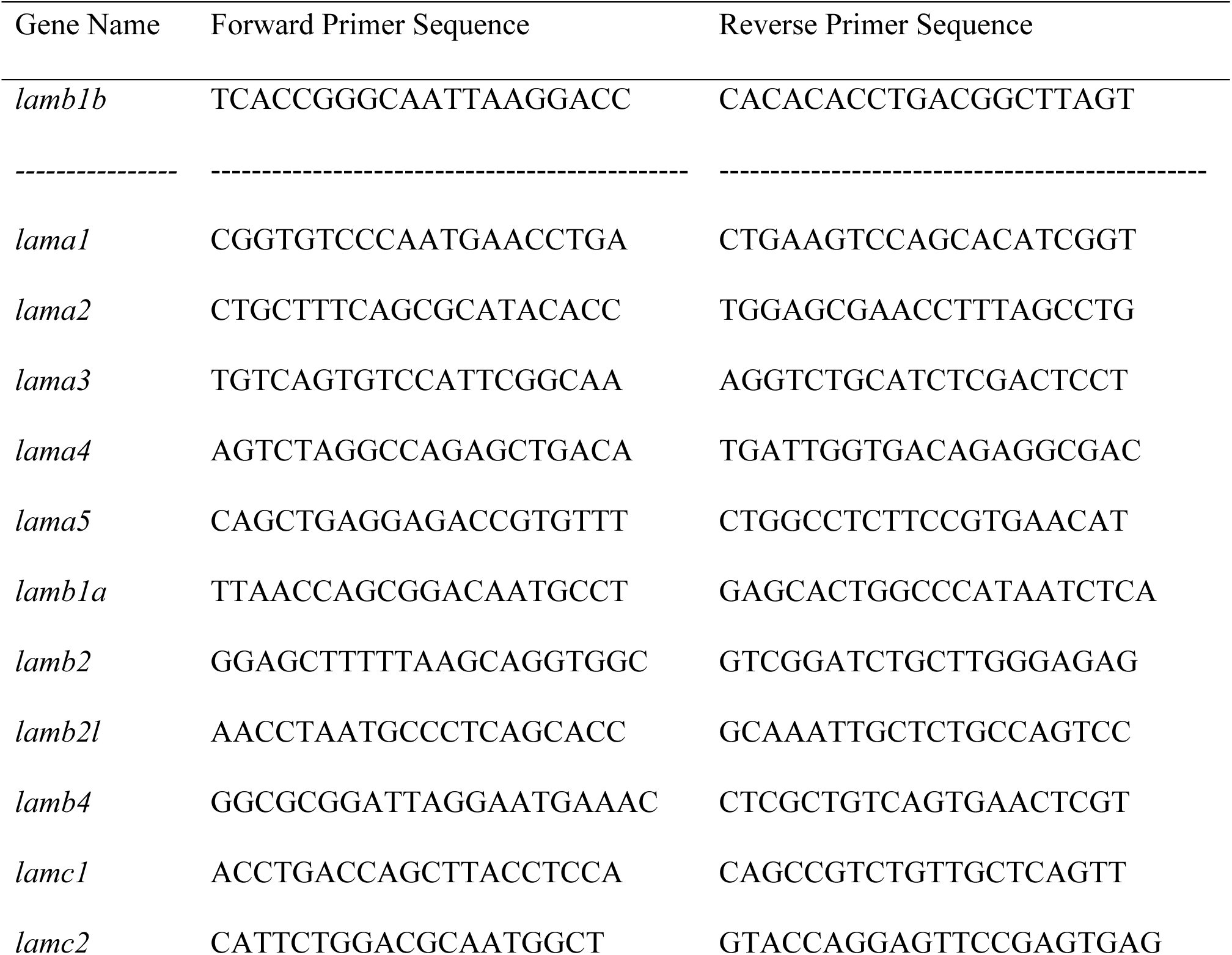
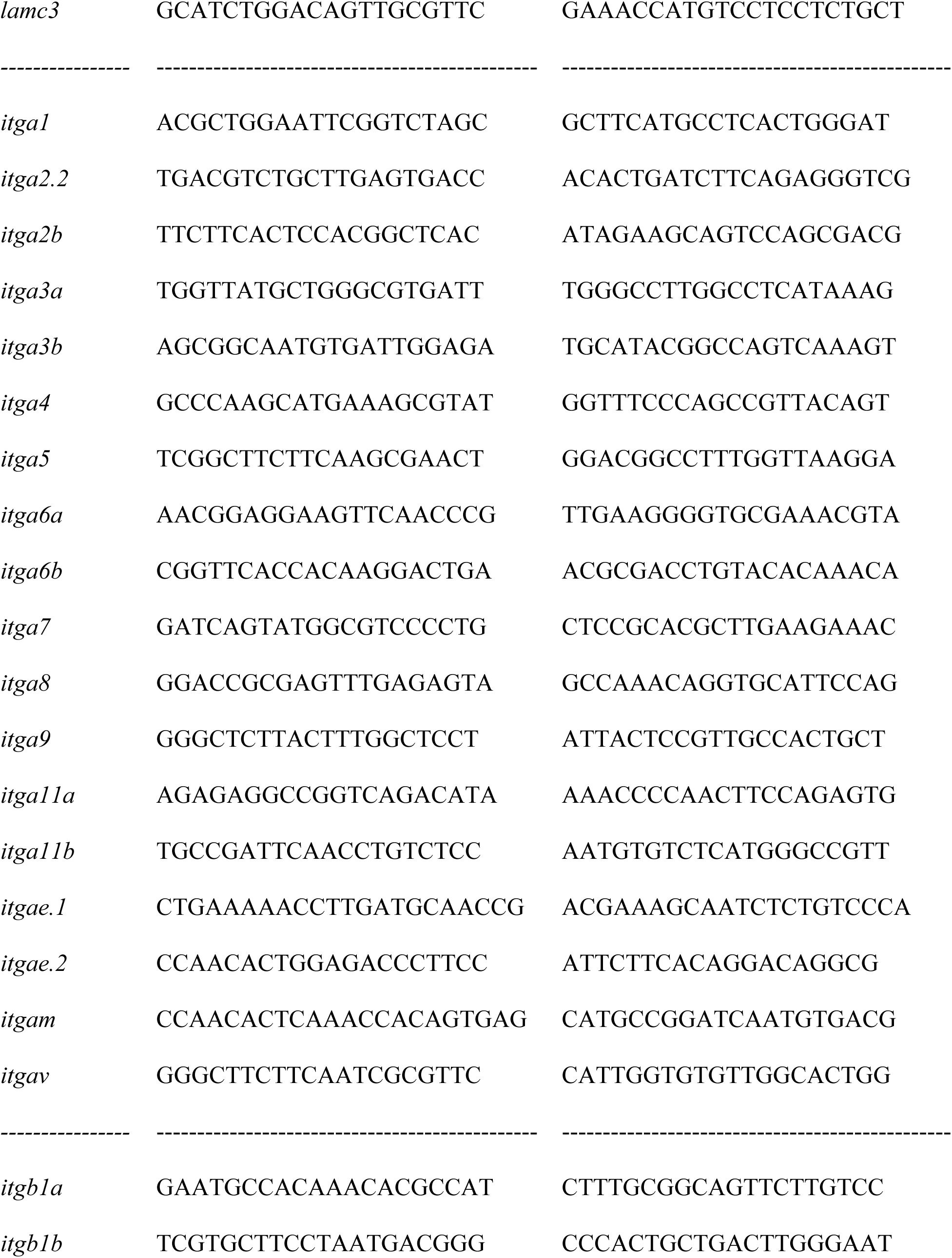
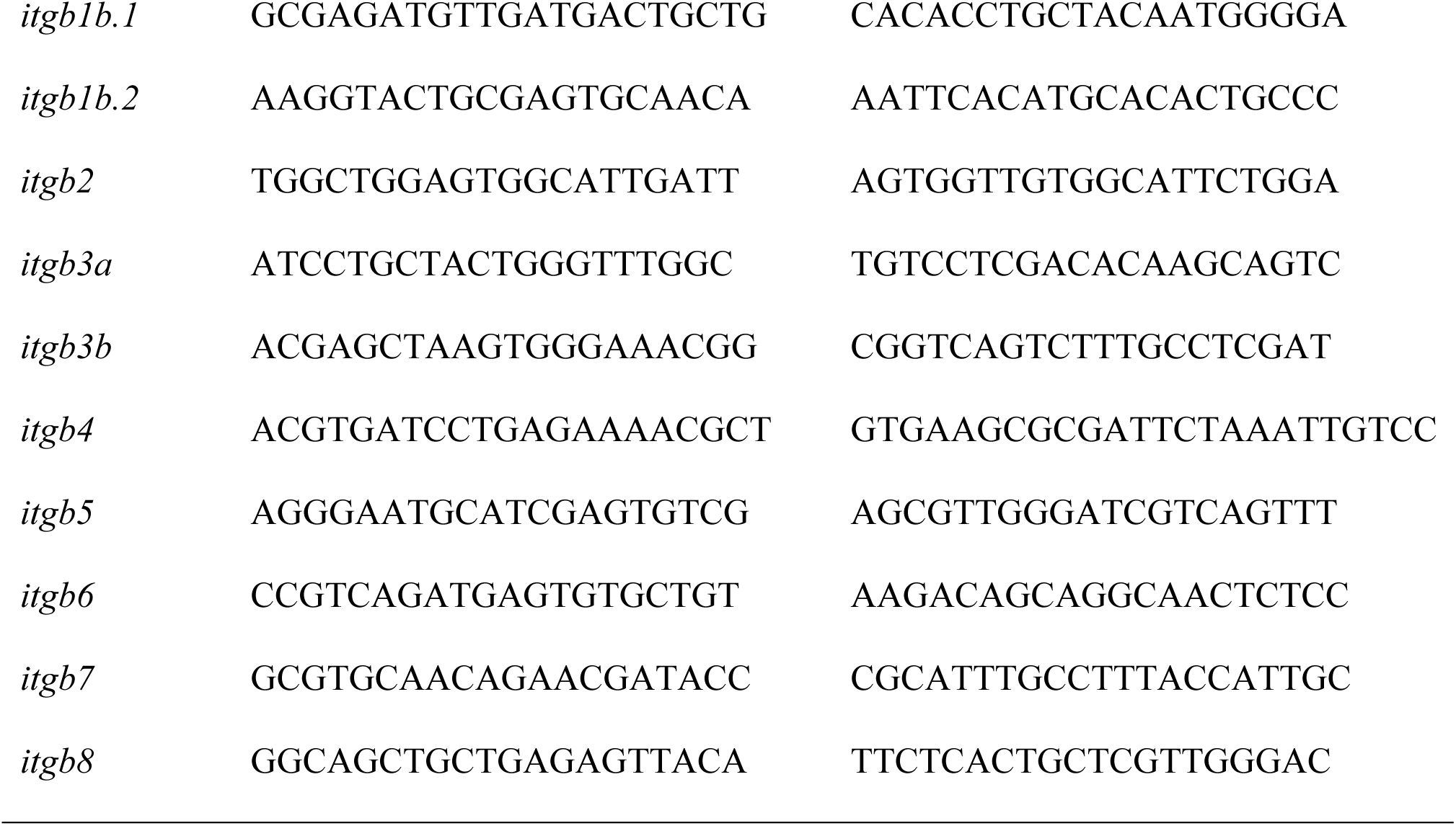
Primer Sequences.

### RNA fluorescent *in situ* hybridization

RNA fluorescent *in situ* hybridization (RNA-FISH) was carried out following the standard protocol for multiplexed hybridized chain reaction (HCR) RNA-FISH provided by Molecular Instruments. Briefly, retinal sections were prepared by immersing slides in 4% PFA for 15 minutes at 4°C, followed by washes in increasing ethanol concentrations for 5 minutes each at RT and incubated in a 10µg/ml proteinase K solution for 10 minutes at 37°C. The slides were washed in 1xPBS for 5 minutes at RT. During the detection stage, molecular probes specific to *lamb1b* transcript were hybridized to the tissue during an overnight incubation period at 37°C. Excess probes were removed by incubating slides at 37°C in 75% probe wash buffer (PWB)/25% 5xSSCT (sodium chloride sodium citrate with 0.1% Tween 20) for 15 minutes, followed by 50% PWB/50% 5xSSCT for 15 minutes, 25% PWB/75% 5xSSCT for 15 minutes, and finally, 100% 5xSSCT for 15 minutes. Finally, during the amplification stage, metastable fluorescent HCR hairpins specific to the type of probe used during the previous step were incubated overnight on the tissue at RT to detect the probes and amplify the fluorescent signal.

## RESULTS

### *lamb1b* gene is expressed in the Müller glia in damaged and undamaged retinas

Given the crucial role β1-containing laminins play in retinal development (Halfter et al., 2008; Edwards and Lefebvre, 2013; Varshney et al., 2015), we examined the expression of *lamb1* genes in the regenerating retina using qRT-PCR. Zebrafish have two *lamb1* isoforms – *lamb1a* and *lamb1b*, however only *lamb1b* is significantly upregulated in the regenerating retina. Initially, *lamb1b* expression is downregulated at 36h of constant light, which corresponds to when the Müller glia (MG) begin to proliferate (Figure 1A). This is followed by a large increase in *lamb1b* expression at 48h of light damage, which coincides with the completion of the MG asymmetric cell division and re-entry of the Müller glia-derived Progenitor Cells (MGPCs) into the cell cycle (Figure 1A).

**Figure 1.**
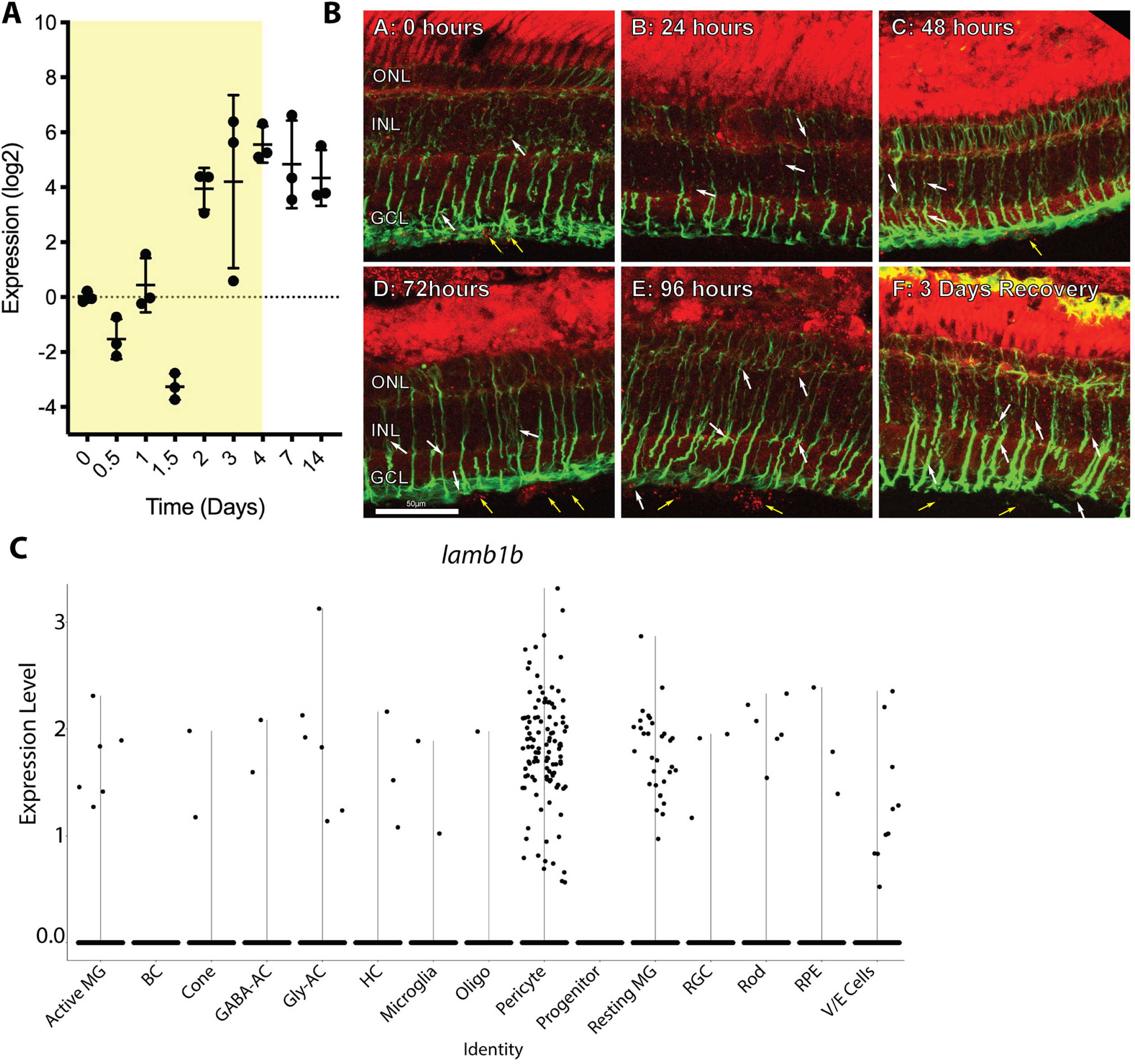
*Lamb1b* is expressed by Müller glia. **A:** qRT-PCR analysis of *lamb1b* gene expression in the zebrafish retina throughout the light damage (yellow region) and recovery paradigm. The expression is downregulated at 36h, and increases from 48h onward, correlating with MG mitosis. **B:** Adult *albino* zebrafish were dark adapted for 14 days and subjected to intense light treatment for 0, 24, 48, 72, and 96 hours. Additionally, some fish were subjected to 96 hours of light damage and allowed 3 days of recovery before tissue harvest. *lamb1b* expression, labeled by RNA-FISH, is shown by the red puncta and GFAP immunolabeling is shown in green. The results indicate that *lamb1b* is expressed diffusely throughout the undamaged and regenerating retinas. *lamb1b* transcripts are observed in Müller glia cell bodies and processes (yellow puncta), particularly in the regenerating retinas (**white arrows**), and the vasculature. **C:** Violin plot from light-damaged snRNA-Seq dataset (Hoang et al., 2000) confirms that *lamb1b* expression is restricted to a small percentage of Müller glia and pericytes in the adult retina.

Next, we examined the spatial and temporal expression of *lamb1b* using HCR-FISH on retinal cryosections at 0, 24, 36, 48, 72, and 96 hours of light damage. The sections were colabeled with anti-GFAP antibodies to visualize the MG (Figure 1B). In the undamaged retina, the *lamb1b* probe primarily labeled the MG endfeet and processes, the vasculature region, and at a low level in the ONL (Figure 1B). These findings are consistent with previous reports of laminin expression by the MG and photoreceptors (Libby et al., 2000b). Upon light damage, however, we observed a loss of the MG endfoot signal localization, and its redistribution throughout the MG processes (Figure 1B). This suggests a loss of cell polarity, consistent with gliosis observed in various glia types (Fisher et al., 2005; Robel et al., 2011). Beginning at 48h, the signal relocalizes to the MG endfeet and vasculature, suggesting resumed MG polarity and laminin secretion into the ECM following cytokinesis (Figure 1B). This *lamb1b* expression to the MG is consistent with the MG being the major source of many ECM molecules in the retina (Pouw et al., 2021). To confirm this expression pattern, we examined *lamb1b* expression in the snRNA-Seq dataset from light-damaged adult zebrafish retinas (Hoang et al., 2000). We found that *lamb1b* was expressed primarily in pericytes, resting MG, and activated MG (Figure 1C), which is consistent with the HCR-FISH analysis.

### β1b chain-containing laminins regulate the MG and PNC proliferation in the regenerating retina

We next examined whether Lamβ1-containing laminins play a role in the MG/MGPC proliferation response following light damage. We used morpholino-mediated gene knockdowns to inhibit Lamβ1b translation in the *albino*; *Tg(gfap:EGFP)^nt11^* adult fish retina and assessed proliferation in retinal cryosections throughout the regeneration time course by immunolabeling with anti-PCNA antibodies. The sections were colabeled with anti-GFP antibodies to visualize MG and MGPCs. We used a Standard Control morpholino (SC-MO) that is not complementary to any known sequence in the zebrafish genome (GeneTools, LLC) and a *pcna* morpholino as negative and positive controls, respectively (Figure 2B).

**Figure 2.**
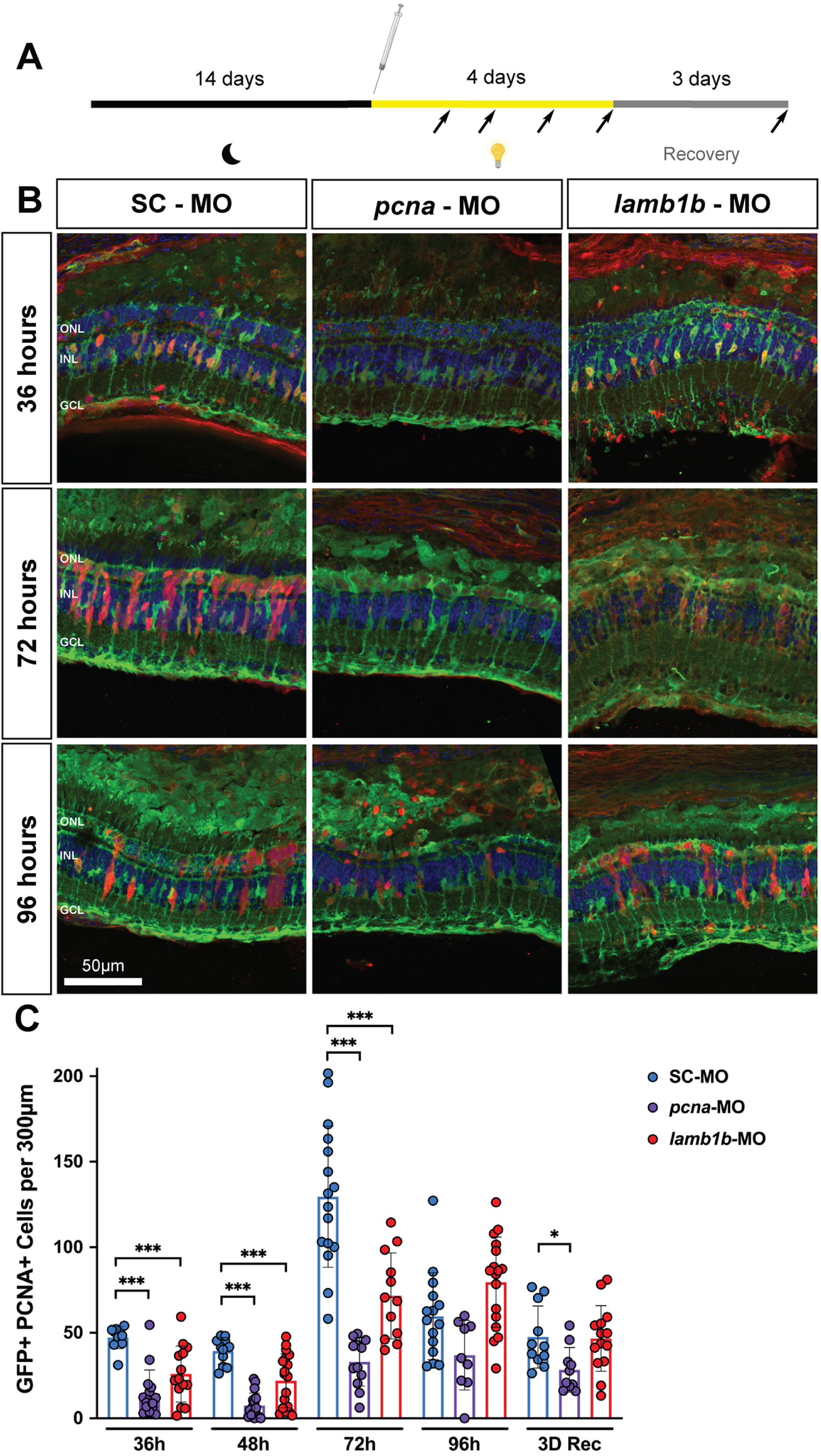
β1b chain-containing laminins regulate regeneration of the zebrafish retina. **A:** Experimental approach involved dark adapting *Tg(gfap:EGFP)* zebrafish for two weeks and then intravitreally injecting and electroporating either the Standard Control (SC-MO), the anti-*pcna* (*pcna*-MO), or anti-*lamb1b* morpholino (*lamb1b* - MO) into the retina. The black arrows indicate tissue collection timepoints. **B:** Cryosections of SC-MO, anti-*pcna,* and anti-*lamb1b* morpholino-injected retinas collected at 36, 72, and 96 hours of light damage. Knocking down *lamb1b* resulted in a reduction of PCNA+ cells in the retina. **C**: Quantification of GFP+ PCNA+ cells (MG and MGPCs) per 300μm of the INL throughout the light damage time course. The *lamb1b* morphants display a significant reduction in the number of proliferating MG and MGPCs at 36, 48, and 72 hours of light damage, suggesting that Lamβ1b regulates the regenerative response in the retina following light damage. * - p≤0.05; ** - p≤0.01; *** - p≤0.001.

We observed a significant reduction in the number of proliferating MG and MGPCs (GFP+ and PCNA+) in the INL at 36, 48, and 72h of light damage with both the *pcna* and *lamb1b* morpholinos relative to the SC-MO retinas, suggesting that β1b-containing laminins are required for MG reprogramming and MGPC proliferation (Figure 2B-C). A second *lamb1b* morpholino (*lamb1b-MO#2*) phenocopies the significant reduction in MG and MGPCs proliferation relative to the SC-MO at 36 and 72h of light damage (Supplemental Figure 1). The effect of the Lamβ1b knockdown on the number of proliferating cells diminishes at 96 hours and 3 days of recovery, suggesting that β1b chain-containing laminins play a role during the initial stages of MG reprogramming, MGPC genesis, and MGPC amplification, which correspond to 36, 48, and 72 hours of light damage, respectively (Gorsuch and Hyde, 2014), but may not play a role in later stages of regeneration. Alternatively, the observed results may be due to the loss of morpholino efficacy due to degradation and derepression of *lamb1b* expression at later time points.

### β1b-containing laminins regulate cell survival in the ONL

Previous studies reported that the number of proliferating cells during regeneration is directly proportional to the amount of cell death in the damaged retina (Montgomery et al., 2010). Therefore, we examined the effect of the Lamβ1b knockdown on retinal cell death using the TUNEL assay at 12, 24, 36, and 48h of light damage. Consistent with previous reports (Gorsuch and Hyde, 2014), we observed peak photoreceptor apoptosis at 24h of light damage. A significant increase in the number of TUNEL+ ONL cells was observed in *lamb1b* morphants relative to SC-MO control retinas at 24h of light damage (Figure 3B-C), but not at the other timepoints. These data suggest that β1b chain-containing laminins may serve a mild neuroprotective role, given that a reduction of Lamβ1b expression increases cell death at 24 hours. However, a neuroprotective effect would not explain the reduction in the number of proliferating MG and MGPCs that we observed in the *lamb1b* morphants during regeneration (Figure 2C). Therefore, it is likely that the influence of β1b chain-containing laminins on the regenerative response of the MG is not a consequence of suppressing apoptosis pathways.

**Figure 3.**
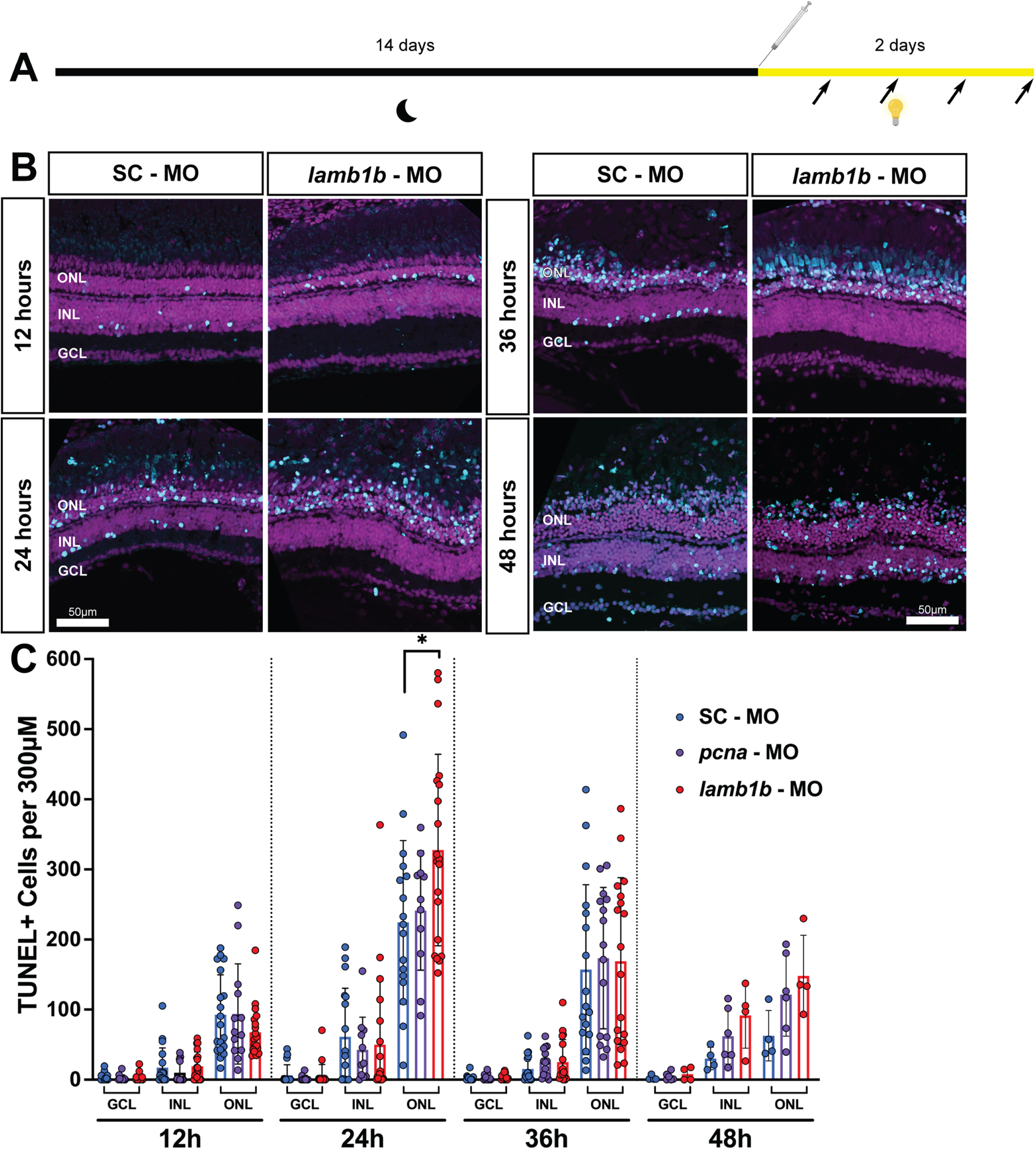
β1b chain-containing laminins regulate cell survival in the outer nuclear layer. **A:** Experimental approach. Adult *albino*; *Tg(gfap:EGFP)* zebrafish were dark-adapted for two weeks before treatment with either Standard Control, anti-*pcna*, or anti-*lamb1b* morpholinos (represented by the Hamilton syringe). The fish were then moved into intense constant light for either 12, 24, 36, or 48 hours before tissue collection. **B:** Cryosections of Standard Control morpholino and anti-*lamb1b* morpholino-injected retinas at 12, 24, 36, and 48 hours of light damage. DAPI (magenta) labels nuclei. Apoptotic cells detected by the TUNEL assay are labeled in cyan. **C:** Quantification of TUNEL+ cells per 300μm at 12, 24, 36, and 48 hours of light damage in the ganglion cell layer (GCL), inner nuclear layer (INL), and the outer nuclear layer (ONL) in eyes treated with either Standard Control, anti-*pcna*, or anti-*lamb1b* morpholinos. The Lamβ1b knockdown results in a significant increase of apoptotic cells within the ONL at 24 hours, suggesting that more photoreceptors are dying relative to the control. * - p≤0.05.

### β1b chain-containing laminins do not influence MGPC cell fate in the regenerating retina

Several studies demonstrated that laminins are involved in cell fate determination in the developing retina. It was previously shown that laminin-111, which is composed of *α*1, β1, and *γ*1 chains, influenced a retinoblastoma-derived cell line to differentiate *in vitro* into either cells with neuronal-like characteristics or photoreceptors, depending on the presence or absence of laminin-111, respectively (Campbell and Chader, 1988). Because regeneration has many similarities to retinal development (Lahne et al., 2021; Lyu et al., 2023), we investigated the role of β1b chain-containing laminins in cell-fate determination in the regenerating zebrafish retina. We intraperitoneally injected EdU at 72 hours and 96 hours of light damage, which correspond to peak MGPC proliferation and migration during light-induced retinal damage and regeneration, as well as at 1 day of recovery (1D Rec) to assess the late-stage neurogenesis in the regenerating retina. Tissues were collected for analysis after 7 days of recovery. Consistent with the proliferation results (Figure 2C), we observed a significant reduction in the number of EdU+ cells in 72h samples in the *lamb1b* morphants relative to the Standard Control morphant (Figure 4B).

**Figure 4.**
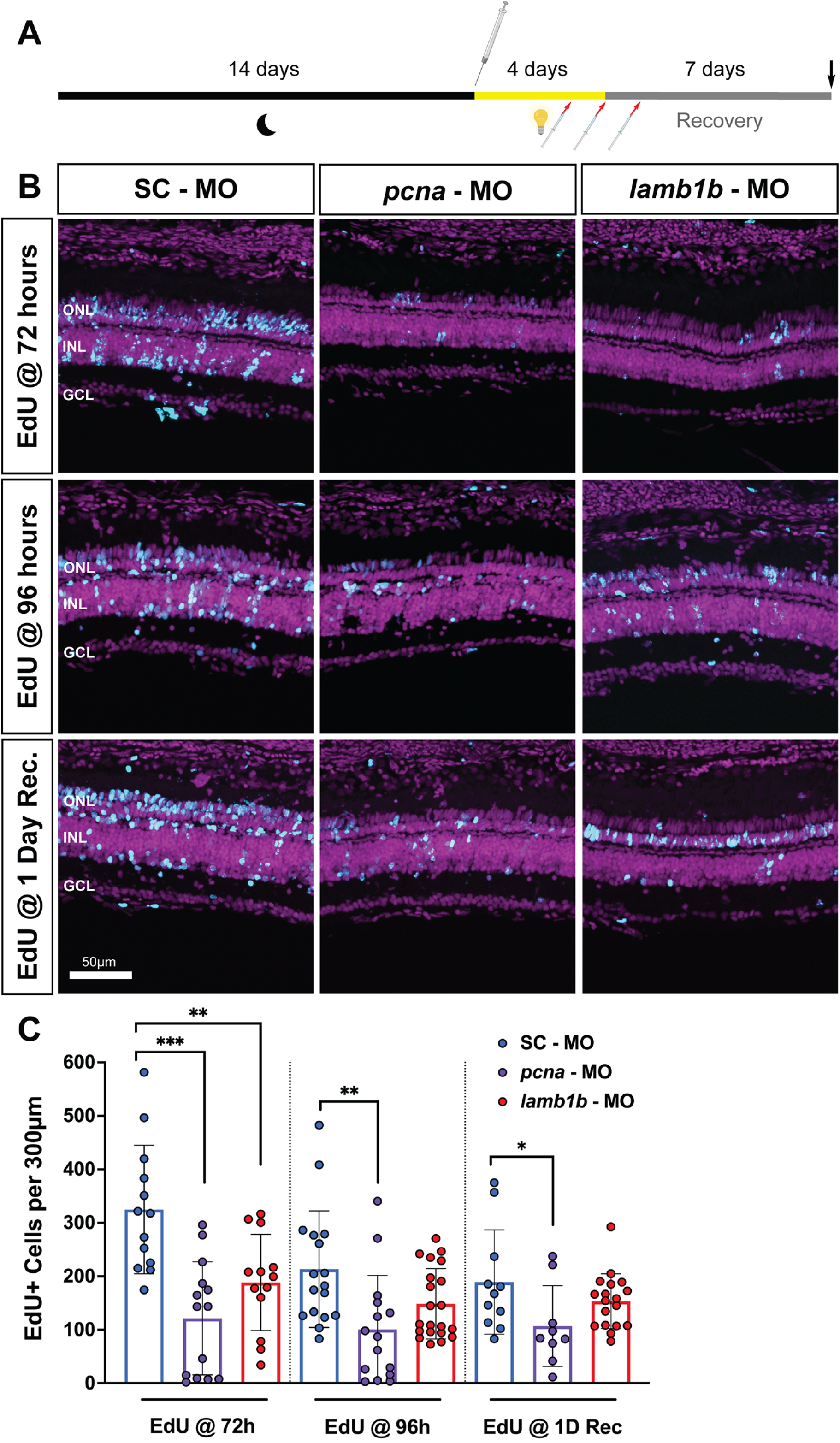
Reduced numbers of EdU+ cells in *Lamb1b* morphant retinas following 7 days after recovering from light damage. **A:** Experimental approach. Adult *albino* zebrafish were dark-adapted for two weeks, followed by electroporation with either Standard Control, anti-*pcna*, or anti-*lamb1b* morpholinos (represented by the Hamilton syringe). Subsequently, the fish were exposed to intense constant light. Fish were divided into 3 groups, receiving a single EdU injection at 72 or 96h of light damage, or at 1-day recovery. The red arrows/syringes represent these EdU injection times. At four days of light damage, the fish were removed from constant light treatment and placed in a normal light:dark cycle (recovery) for 7 days until tissue collection (black arrow). **B**: Retinal cryosections collected at 7 days of recovery, which were treated with a single EdU pulse at 72h, or 96h, or 1DR. **C:** Quantification of the overall numbers of EdU+ cells per 300μm within experimental retinas. There are significantly fewer EdU+ cells in the Lamβ1b knockdown relative to the samples treated with Standard Control morpholinos labeled at 72h light. * - p≤0.05; ** - p≤0.01; *** - p≤0.001.

In addition to quantifying the number of EdU+ cells, we colabeled the cryosections with cell-specific markers for ganglion/amacrine cells (HuC/D), bipolar cells (PKC*α*), Müller glia (GFAP), and rod photoreceptors (4c12). The proportion of EdU+ cells that colabeled with each cell-specific marker was quantified to determine the relative proportions of cell types that differentiated from the MGPCs during regeneration. Cone photoreceptors were identified based on morphology as oblong nuclei apical of the ONL, without the use of a specific antibody. We observed that the differences in cell populations produced during light-induced regeneration between the *lamb1b* morphants and the Standard Control retinas were mostly not significant (Figure 5), indicating that the morpholino-mediated knockdown of Lamβ1b does not influence cell-fate determination during regeneration. However, we observed that the fraction of regenerated cones was greatest when the EdU was injected at either 72 or 96 hours and then dropped at the 1D recovery timepoint (Figure 5K). In contrast, the fraction of regenerated rods was lowest at the first two EdU injection timepoints and increased at 1D recovery (Figure 5I). This suggests that MGPCs commit to becoming cones before committing to becoming rods during regeneration of the light-damaged retina. Further, we observed an unexpected localization of HuC/D+ cells within the apical portion of the INL and in the ONL of Standard Control and *lamb1b* morphants following regeneration. The presence of these ectopic cells highlights the difference between regeneration and development and underlines the fact that regeneration of the zebrafish retina does not result in the same cellular makeup of the tissue as that of the undamaged tissue.

**Figure 5.**
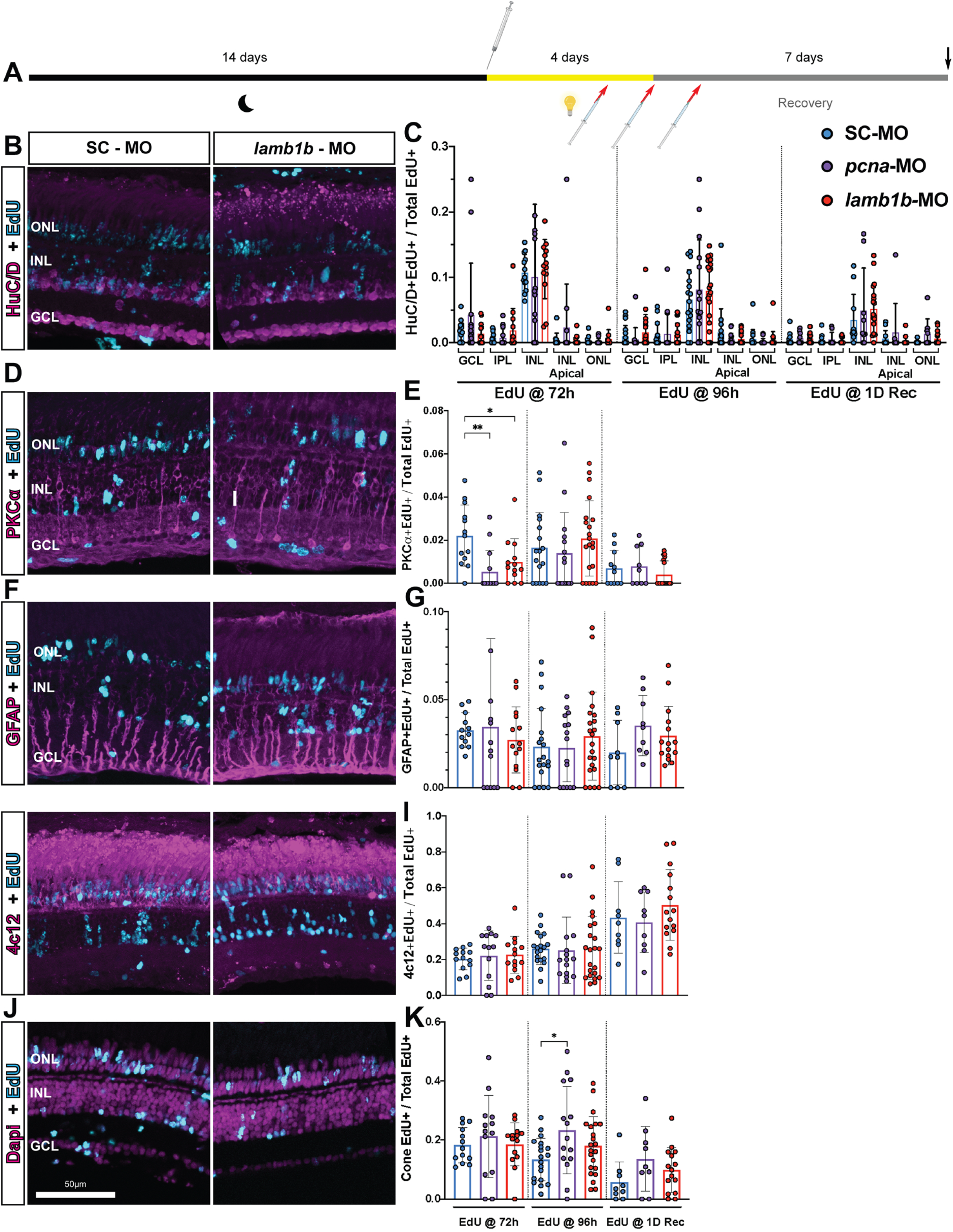
β1b chain-containing laminins do not appear to significantly influence MGPC cell fate in the regenerating retina. **A:** Experimental approach. Adult *albino* zebrafish were dark-adapted for two weeks prior to being treated with either Standard Control, anti-*pcna,* or anti-*lamb1b* morpholinos (denoted by the Hamilton syringe). The fish were then moved into intense constant light to induce retinal injury. The fish were divided into three groups, which received a single pulse of EdU via intraperitoneal injection at either 72 hours of light treatment, 96 hours of light treatment, or 1 day of recovery after light treatment (red arrow/syringe represent EdU injections). The fish were then sacrificed after 7 days of recovery in normal light (represented by the black arrow). **B:** Retinal cryosections collected at 7 days of recovery from fish receiving an EdU injection at 72 hours, colabeled with HuC/D (ganglion and amacrine cells). **C:** Quantification of EdU+HuC/D+ cells per 300μm in recovered retinas. The analysis was divided by cell layers due to presence of ectopic HuC/D+ cells in the tissue following recovery. **D:** Retinal cryosections collected at 7 days of recovery from fish receiving an EdU injection at 72 hours, colabeled with PKCα (bipolar cells). **E:** Quantification of EdU+PKCα+ cells in the regenerated retinas. **F:** Retinal cryosections collected at 7 days of recovery from fish receiving an EdU injection at 72 hours, colabeled with GFAP (Müller glia). **G:** Quantification of EdU+GFAP+ cells per 300μm in the regenerated retinas. **H:** Retinal cryosections collected at 7 days of recovery from fish receiving an EdU injection at 72 hours, colabeled with 4c12 (rod photoreceptors). **I:** Quantification of EdU+4c12+ cells per 300μm in the regenerated retinas. **J:** Retinal cryosections collected at 7 days of recovery from fish receiving an EdU injection at 72 hours. **K:** Quantification of EdU+ cone photoreceptors per 300μm in the regenerated retinas. Cones were assessed morphologically, as oblong nuclei apical of the ONL, which contains rods.

At the same time, we observed a statistically significant decrease in the proportion of EdU+ PKC*α*+ (bipolar) cells in the *lamb1b* morphants relative to the control retinas when EdU was administered at 72 hours of light damage, the peak of MGPC proliferation. In control retinas, 2.21% of EdU+ cells were also PKC*α*+ relative to 0.999% of EdU+ cells that colabeled with PKC*α*+ in *lamb1b* morphants (p<0.05). Additionally, when we analyzed similar retinas after 28 days of recovery, there were no significant differences between 7 and 28 days of recovery (data not shown). These findings are consistent with our previous work, which demonstrates there is not significant pruning of newly generated retinal cells following regeneration (Lyu et al., 2023).

### β1b chain-containing laminins regulate the expression of other laminins and their cellular receptors

To begin elucidating the molecular mechanism through which β1b chain-containing laminins regulate Müller glia during retinal regeneration, we investigated the effect of the Lamβ1b knockdown on the expression of integrins, the primary cell-surface receptor for laminin molecules. qRT-PCR was used to quantify the expression of several integrin chain genes in the *lamb1b* morphants, relative to the Standard Control, at 72 hours of light damage. Since laminins were previously shown to regulate the special localization of integrins in the retina (Serjanov et al., 2018) and are known to regulate their own receptor expression (Condic and Letourneau, 1997), we investigated integrin expression in the morphant tissues. Indeed, knockdown of Lamβ1b expression results in altered expression of various integrin-family receptors, which mediate binding between laminins and the retinal cells (Figure 6A). In the *lamb1b* morphants, expression of *itga2.2*, *itga3a*, *itga3b*, *itga4*, *itga5*, *itga11a*, and *itgb4* were significantly reduced relative to the Standard Control retinas. In contrast, *itga2b* expression was upregulated in the *lamb1b* morphants (Figure 6A).

**Figure 6.**
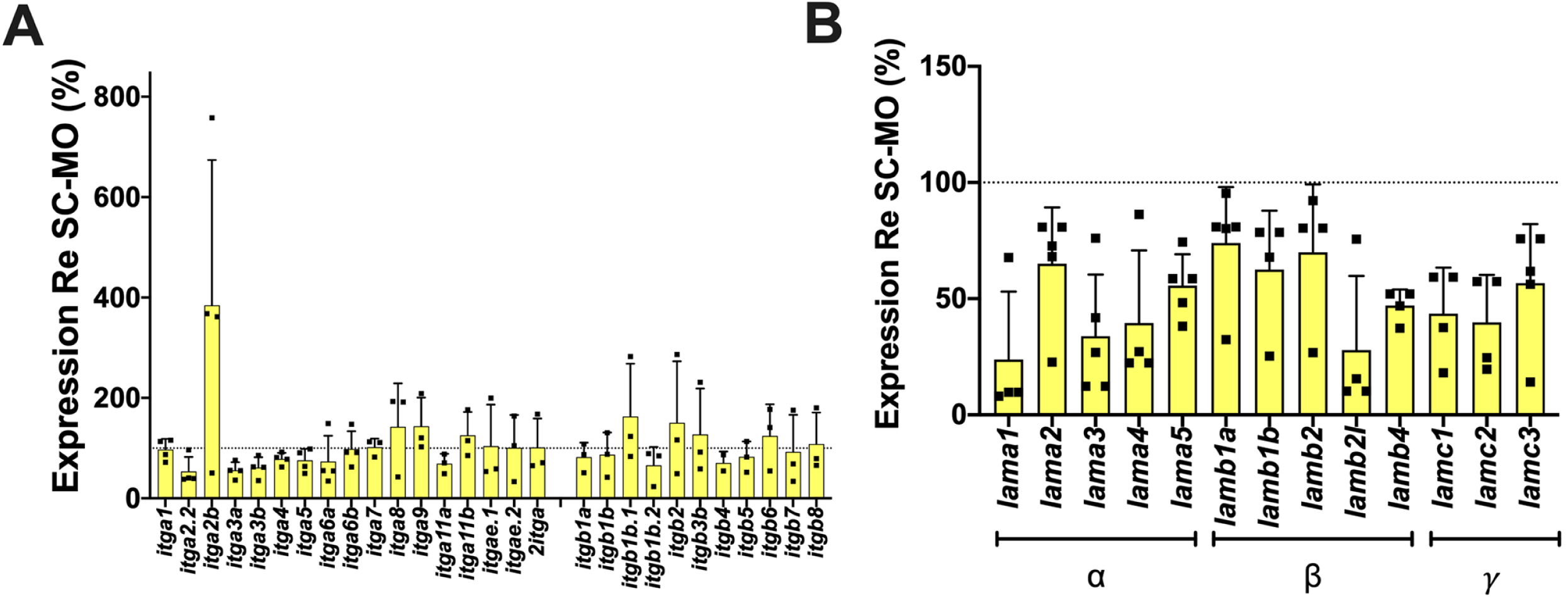
β1b chain-containing laminins regulate expression of integrin receptors and other laminin genes. **A**: The knockdown of Lamβ1b results in altered expression of various integrin-family receptors, which mediate binding between laminins and the retinal cells. **B:** The knockdown of Lamβ1b expression results in the downregulation of other laminin chains. These data indicate that the *lamb1b* morphant has broad effects on the expression of other laminin chains.

Because the laminin chains must function as a heterotrimer, with the β1 chain combining with alpha and gamma subunits, it is possible that Lamβ1b knockdown may also affect the expression of other laminin subunits. Therefore, we also analyzed laminin gene expression in the *lamb1b* morphant at 72h of light damage relative to Standard Control morphant retinas. Interestingly, most other laminin chains decrease in expression in the *lamb1b* morphants, suggesting that the inhibition of Lamβ1b translation influences the expression levels of other laminin chains (Figure 6B).

## DISCUSSION

While several studies examined the role of the extracellular matrix (ECM) during retinal development (Varshney et al., 2015), it was previously unclear if the ECM is also involved in zebrafish retinal regeneration. Laminins, which are major components of ECM, are known to influence photoreceptor cell-fate decisions, morphogenesis of rod inner and outer segments, as well as synaptogenesis during vertebrate retinal development (Libby et al., 2000a). While neuronal regeneration in the zebrafish retina is not a recapitulation of developmental programs (Lyu et al., 2023), there are similar processes, such as cellular migration, apoptosis, and differentiation (Lahne et al., 2020, 2021). Therefore, it is likely that ECM components, such as laminins, will also serve a necessary function during regeneration, and a greater understanding of the role of the ECM during retinal regeneration in zebrafish may reveal how the ECM promotes self-healing pathways in the zebrafish eye while prohibiting regeneration in mammals.

In this study, we examined the role of the laminin β1 chain, which is a necessary component of all β1 chain-containing laminin molecules, in regeneration of the zebrafish retina. At the outset, we believed that Lamβ1b was involved in retinal regeneration because *lamb1b* expression significantly increased during light-induced retinal injury and regeneration in adult zebrafish (Figure 1A). RNA *in situ* hybridization confirmed that *lamb1b* is expressed at low levels in the MG of undamaged retinas and potentially by other cell types, such as bipolar cells and photoreceptors, as we observe its expression in both the INL and ONL. Further, *lamb1b* expression increased in the MG during retinal regeneration. Based on the temporal and spatial *lamb1b* expression in the light-damaged zebrafish retina, we examined whether Lamβ1b, is required during retinal regeneration in zebrafish. We found that *lamb1b* morphants exhibited fewer proliferating MG and Müller glia-derived Progenitor Cells (MGPCs) between 36 and 72 hours of light damage, suggesting that Lamβ1b-containing laminins are necessary for MG proliferation in the regenerating zebrafish retina. It was previously shown that MG reprogramming and cell cycle reentry occurs by 36 hours of light damage (Gorsuch and Hyde, 2014), which is consistent with β1b chain-containing laminins (β1b CCLs) being required for MG reprogramming and/or cell cycle reentry. It is possible that β1b CCLs are necessary for providing external signals that facilitate the regenerative capabilities of the MG, similar to TNFα being produced by dying retinal neurons to stimulate MG proliferation (Nelson et al., 2013; Conner et al., 2014). Alternatively, β1b CCLs may bind and release a regeneration signal.

It is believed that the number of proliferating MG during retinal regeneration is directly proportional to the amount of cell death (Montgomery et al., 2010). Therefore, we assessed whether the reduced proliferation observed in the *lamb1b* morphants resulted from a reduced number of apoptotic cells. We found that the number of TUNEL+ cells in the *lamb1b* morphants did not differ significantly from the controls within retinal INL and GCL layers at various timepoints throughout light damage. Interestingly, we did observe an increased number of ONL TUNEL+ cells in the *lamb1b* morphants relative to the controls at 24 hours of light damage, which corresponds to the time of maximal cell death in the light-damaged retina (Fig. 3C; (Gorsuch and Hyde, 2014). This suggests that β1b CCLs may serve a neuroprotective role for the photoreceptors in light-damaged retinas. It has been established that ECM components of the inner photoreceptor matrix (IPM), which surrounds photoreceptors in the ONL, are involved in regulating photoreceptor survival and cell death by controlling the concentrations of soluble molecules, such as FGF-2 (Faktorovich et al., 1990; Carwille et al., 1998; Libby et al., 2000a). While β2 CCLs are the predominant laminin found in the mammalian IPM (Libby et al., 2000b), it is possible the β1b CCLs may serve a similar role in fish, given that we observed *lamb1b* transcripts in the ONL. The presence of β1b CCLs in the IPM would be consistent with these laminin molecules serving a neuroprotective role, promoting photoreceptor survival either through direct signaling or by regulating the concentration of soluble molecules. Importantly, an increased number of TUNEL+ cells in the *lamb1b* morphants cannot explain fewer proliferating MG that we observed in previous experiments. Therefore, β1b CCLs must regulate retinal regeneration through another mechanism that is unrelated to cell death pathways.

Given that laminins influence cell fate decisions during retinal development (Campbell and Chader, 1988; Hunter et al., 1992; Hunter and Brunken, 1997), we hypothesized that β1b CCLs may serve a similar role during retinal regeneration. A potential mechanism for this process is that physical contact between retinal progenitor cells and ECM components of the basement membrane (inner limiting membrane of the retina) facilitates ECM-cell signaling to modulate mitotic spindle orientation and cell cycle dynamics, which influence cell fate decisions (Serjanov et al., 2018, 2022). However, we did not observe significant differences in the cell fates adopted by MGPCs during retinal regeneration in the *lamb1b* morphants relative to the control retinas. We conclude from these data that although β1b CCLs are involved in and required for MG and/or MGPC proliferation, they do not have a significant impact on cell fate determinations during retinal regeneration.

One possible exception of the β1b CCLs not altering cell fate determination during retinal regeneration is the apparent effect on bipolar cell regeneration. Previous reports demonstrated that exposing retinal cultures to β2-rich laminin matrices resulted in a six-fold increase in the number of rods, and a three-fold decrease in the number of bipolar cells (Hunter and Brunken, 1997). We did observe a two-fold decrease in the number of newly generated bipolar cells in *lamb1b* morphants relative to Standard Control morphants (Figure 6D-E). These data support the importance of laminins in retinal cell type genesis and serve as additional evidence in the growing body of literature that demonstrates the role of ECM in regulating retinal cell biology.

Finally, we examined the effect of the Lamβ1b knockdown on the expression of integrin receptors, one of the major cell receptors that bind ECM proteins. It was shown that laminin molecules regulate the expression of their own receptors (Condic and Letourneau, 1997). Based on our findings, we aimed to identify integrin receptors that were differentially expressed in response to binding β1b CCLs. We observed that the *lamb1b* morphants broadly affected the expression of integrin chain genes relative to controls. These data provide a starting point for future investigations, which could involve knocking down integrin chains of interest and observing the effect on MG and MGPC proliferation during retinal regeneration. Additionally, we measured the expression changes of other laminin chains in the presence of the Lamβ1b knockdown, finding that most other laminin chains are downregulated in the *lamb1b* morphants relative to the controls. One explanation for this is that β1b CCLs regulate the expression of other laminin chains. Given this effect on the expression of other laminins, it is possible that the phenotypes we observed are not directly attributable to β1b CCLs, but instead to the effect of β1b CCLs on the expression of other laminin chains, which together modulate the composition of the basement membrane. Although the direct or indirect mechanism by which β1b CCLs influence MG behavior is still under investigation, we have shown that β1b CCLs are a necessary component of the ECM during retinal regeneration.

## ACKNOWLEDGEMENTS

This work was supported by a grant from the National Eye Institute-National Institutes of Health to D.R.H. (R01-EY034493-01), the Hiller Family Endowment for Excellence in Stem Cell Research (J.S.), and the Centers for Zebrafish Research and the Center for Stem Cells and Regenerative Medicine at the University of Notre Dame. We thank the Freimann Life Sciences Center technicians for outstanding care and husbandry of the zebrafish, the University of Notre Dame Integrated Imaging Facility and the Optical Microscopy Core for assistance with confocal imaging, and members of the Hyde lab for thoughtful discussions.

---

**Supplemental Figure 1. A second *lamb1b* morpholino phenocopies the suppression of MG and MGPC proliferation.**

**A:** *Tg(gfap:EGFP)* zebrafish that were dark adapted for two weeks, were intravitreally injected and electroporating with either the Standard Control (SC-MO), the anti-*pcna* (*pcna*-MO), or anti-*lamb1b* morpholino (*lamb1b* – MO#2) into the retina. Cryosections of SC-MO, anti-*pcna,* and anti-*lamb1b-MO#2* injected retinas collected at 36 and 72 hours of light damage. Knocking down *lamb1b* resulted in a reduction in the number of PCNA+ cells in the retina. **B**: Quantification of GFP+ PCNA+ cells (MG and MGPCs) per 300μm of the INL (36 and 72 h light) and ONL (72h light). The *lamb1b* morphants display a significant reduction in the number of proliferating MG and MGPCs at both 36 and 72 hours of light damage relative to the SC-MO, confirming that Lamβ1b regulates the regenerative response in the retina following light damage. ** - p≤0.01; **** - p≤0.0001.

