## Supplementary figures and images for "Regulation of Adult Zebrafish Retinal Regeneration by Lamβ1b-Chain-Containing Laminins"

### Supplemental Figure 1

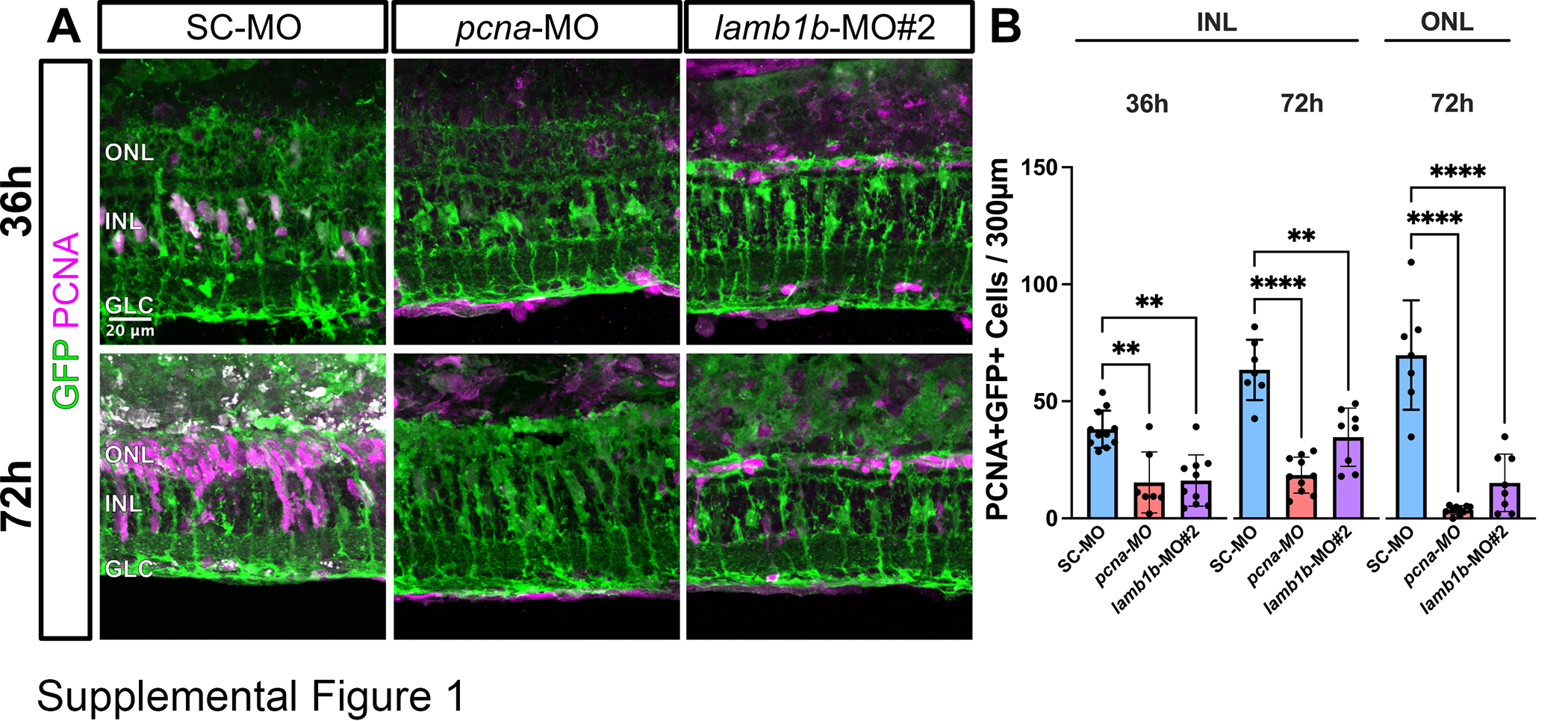
